# Collective Learning in Living Neural Networks Facilitated by Contextual Background Photostimulation

**DOI:** 10.1101/2025.03.03.641104

**Authors:** Dulara De Zoysa, Sylvester J. Gates, Anna M. Emenheiser, Kate M. O’Neill, Wolfgang Losert

**Affiliations:** Institute for Physical Science and Technology, University of Maryland, College Park, Maryland, USA; Fischell Department of Bioengineering, University of Maryland, College Park, Maryland, USA; A. James Clark School of Engineering, University of Maryland, College Park, Maryland, USA

**Keywords:** Spike-timing-dependent-plasticity, Hebbian learning, contextual stimulation, optogenetic stimulation, collective learning

## Abstract

This study explores collective learning in living neural networks, focusing on group-to-group Hebbian learning, i.e., strengthening and weakening of links dependent on the precise timing of their activities. While neuronal plasticity is now well understood for single pairs of neurons, recent research has demonstrated that groups of tens of neurons are required to encode information in mammalian brains. Thus, it is critical to understand how mechanisms of plasticity, in particular spike-timing-dependent plasticity (STDP) operate at the group scale. We find that neuronal groups can reach significant plasticity after only 45 stimuli when a proper tradeoff between pulse duration and photostimulation effectiveness is chosen. Background stimulation, which enhances the reliability of response for the targeted neuronal groups, is necessary for rapid network-level Hebbian learning. By demonstrating enhanced learning in the presence of background activity, this study underscores the highly cooperative character of neurons and the importance of investigating learning, information flow, and memory formation at the network scale.

## I. INTRODUCTION

Spike-timing-dependent plasticity (STDP) is a biological process that adjusts the connection strength between neurons in the brain based on the relative timing of input and output action potentials (or spikes) of each neuron. This process partially explains the activity-dependent development of neuronal networks, which includes long-term potentiation (LTP) and long-term depression (LTD) of neuronal synapses. If an input spike (pre-synaptic spike) to a particular neuron tends to occur immediately before its output spike (post-synaptic spike), then the input connection to the neuron becomes much stronger [1, 2]. If a pre-synaptic spike tends to occur immediately after a post-synaptic spike, then the input connection is made weaker. STDP is a Hebbian learning rule, as it modifies synaptic strength based on the precise timing of pre and post synaptic spikes, reinforcing connections when presynaptic activity precedes postsynaptic firing [3]. This aligns with the Hebbian principle that “neurons that fire together, wire together,” shaping network plasticity and learning. Furthermore, synapses were strengthened if the pre-synaptic spike occurred 3-20 milliseconds before a post-synaptic spike, while a reversal of the spike timing weakened the synaptic connections [4]. If a neuron does not fire, its synaptic connections to the active neurons are not immediately altered, though unused synapses tend to weaken over time.

Most STDP studies have focused on neuronal pairs, allowing for clear, controlled observations of pre- and post-synaptic spike timing and resulting plasticity. Froemke et al. [5] have demonstrated that STDP varies with dendritic location, showing that distal synapses exhibit broader timing windows for potentiation and depression compared to proximal ones. This finding suggests that synaptic modifications are not uniform across a neuron but depend on spatial and temporal integration within dendrites. At the network level, these variations imply that synaptic plasticity in neuronal groups may involve complex, location-dependent interactions rather than simple pairwise STDP, potentially leading to emergent dynamics influenced by dendritic processing and local synaptic organization.

Living neural networks operate under the influence of neuromodulators like dopamine and acetylcholine, which can alter STDP rules in a context-dependent manner. Neuromodulation can reshape the timing and degree of plasticity, making the direct extrapolation of findings from neuronal pairs to networks problematic [6]. Studies at the pair scale generally do not account for these modulatory influences, meaning that conclusions drawn from these simplified settings may not accurately reflect STDP dynamics of neuronal networks. Recently, global stimulation with 1P (one-photon) prior to targeted 2P (two-photon) stimulation of neurons has been demonstrated to enhance the efficacy and precision of subsequent 2P stimulation, further demonstrating another aspect of context dependence on neuronal network activity [7]. Complementing this previous study, Dalgleish et al. [8] have investigated how pre-training with 1P stimulation can effectively prime neurons, enhancing their responsiveness to subsequent 2P stimulation. This dual approach of providing context with global activation followed by targeted manipulation enables a more comprehensive understanding of how sparse neuronal ensembles contribute to perception and behavior, aligning with the sparse coding hypothesis in neural circuits. The sparse coding hypothesis postulates that neural circuits efficiently represent information by activating a small subset of neurons, typically ranging from 10 to 100 cells [8-10].

However, only studying pairs limits our understanding of how STDP relates to actual learning, which occurs within the context of a functional neural network where neurons engage in complex interactions with many other neurons. Looking at sensory information processing, recent optogenetic studies demonstrated that networks of tens of cells are required to reliably store and transmit sensory information [9, 10]. This raises the question of how plasticity is manifested at the scale of such groups.

Neuronal group plasticity involves studying neurons that receive multiple inputs at different times [2]. Thus, STDP must include considerations of temporal integration, i.e., dendritic computing. Due to such nonlinearities, we expect that coincidental firing patterns across multiple neurons influence plasticity in a way that cannot be captured by pairwise studies alone. Consequently, there is a need for models that incorporate broader temporal and spatial interactions for a more accurate characterization of STDP in neural circuits.

In this study, we systematically investigated the group-to-group collective learning dynamics of neuronal subgroups within a larger network. Specifically, we examined the impact of photostimulation pulse duration and the role of neighboring neurons on learning processes. Our findings reveal that incorporating background stimulation as contextual input significantly enhances the reliability of training neuronal groups, particularly at timescales aligned with spike-timing-dependent plasticity (STDP) based Hebbian learning rules. These cooperative effects of STDP, as elucidated in this research, provide valuable insights into neuronal network development and offer a foundation for more advanced in vivo and in silico studies exploring functional network activity related to learning and memory.

## II. MATERIALS AND METHODS

### A. Primary neuron cell culture and viral transduction

Primary rat embryonic cortical neural cells from Sprague Dawley rats were used for the in-vitro experiments to demonstrate group to group collective learning. Embryos of both sexes were obtained from euthanized pregnant rats at embryonic day of gestation 18 (E18) according to and with the approval of the University of Maryland IACUC protocol (R-JAN-18-05, R-FEB-21-04, R-JAN-24-01). Following the dissection of hippocampi and cortices, the cortices were gently triturated using a fire-polished pipette. These cells were then plated onto culture dishes that had been pre-coated with poly-D-lysine, which promotes neuronal cell adhesion and growth. Subsequently, the cultured cells were maintained in neurobasal media and incubated at 37^0^C temperature and 5% carbon dioxide. After incubation for 3 days, neuronal cells were transduced at 3 multiplicity of infection (MOI) with the bicistronic lentiviral vector, pLV[Exp]-Bsd-SYN1-jGCaMP8s-P2A-ChrimsonR-ST, which provides robust co-expression of the Calcium indicator (jGCaMP8s) and the opsin (stChrimsonR) used for holographic optogenetic stimulation [11]. A full media change was performed after 48 h of transduction. Neurons were incubated at 37^0^C temperature and 5% carbon dioxide, while doing half media changes every other day. In-vitro calcium imaging and holographic optogenetic photostimulation were performed one week after transduction at 10 days in vitro (DIV 10).

### B. Optogenetic stimulation and Calcium imaging workflow

The transduced cells were cultured in neurobasal media and maintained at 37°C with 5% carbon dioxide throughout the experiment. Calcium imaging was performed within a 750×750 µm field of view. Imaging and photostimulation experiments were conducted using a Nikon Eclipse Ti2 microscope integrated with a digital micromirror device (DMD; MIGHTEX Polygon). Calcium imaging was performed using 476 nm light, while optogenetic photostimulation was provided with 555 nm light. The experimental workflow leveraged the advanced capabilities of NeuroART, a recently developed software platform designed for real-time analysis and photostimulation of neuronal networks [12]. NeuroART was utilized to execute photostimulation protocols through a TCP/IP communication link established between the DMD control software and the NeuroART platform, ensuring precise coordination and real-time control during experiments.

Initially, the entire field of view underwent global photostimulation using 20-millisecond pulses repeated every 2 seconds for a total duration of 40 seconds, and responsive cells were recruited for the subsequent photostimulation experiments. From these responsive cells, 10 neurons were chosen and divided into two groups, group 1 (G1) and group 2 (G2), each comprising 5 neurons. The photostimulation experiment was divided into three stages: training, testing 1, and testing 2. During training, G1 neurons were stimulated first, followed by G2 neurons, using 15-millisecond pulses every 2 seconds for a total duration of 90 seconds. During the testing 1 stage, only G1 neurons were stimulated 20 times to assess whether G2 neurons would fire in response, indicating synaptic learning. Subsequently, during the testing 2 stage, only G2 neurons were stimulated 20 times to determine whether G1 neurons remained inactive, indicating unidirectional nature of the learning process.

For each distinct illumination pattern delivered by the DMD for photostimulation, a background stimulation pattern consisting of randomly selected 10% of non-targeted pixels was superimposed onto the original illumination pattern. The influence of this background stimulation, referred to as “context”, is further detailed in section III.A. During the testing phases, pixels corresponding to the non-targeted group were excluded from the background stimulation. The overall photostimulation workflow and timing of delivered photostimulation patterns for training and testing are illustrated in Fig. 1.

**Fig. 1.**
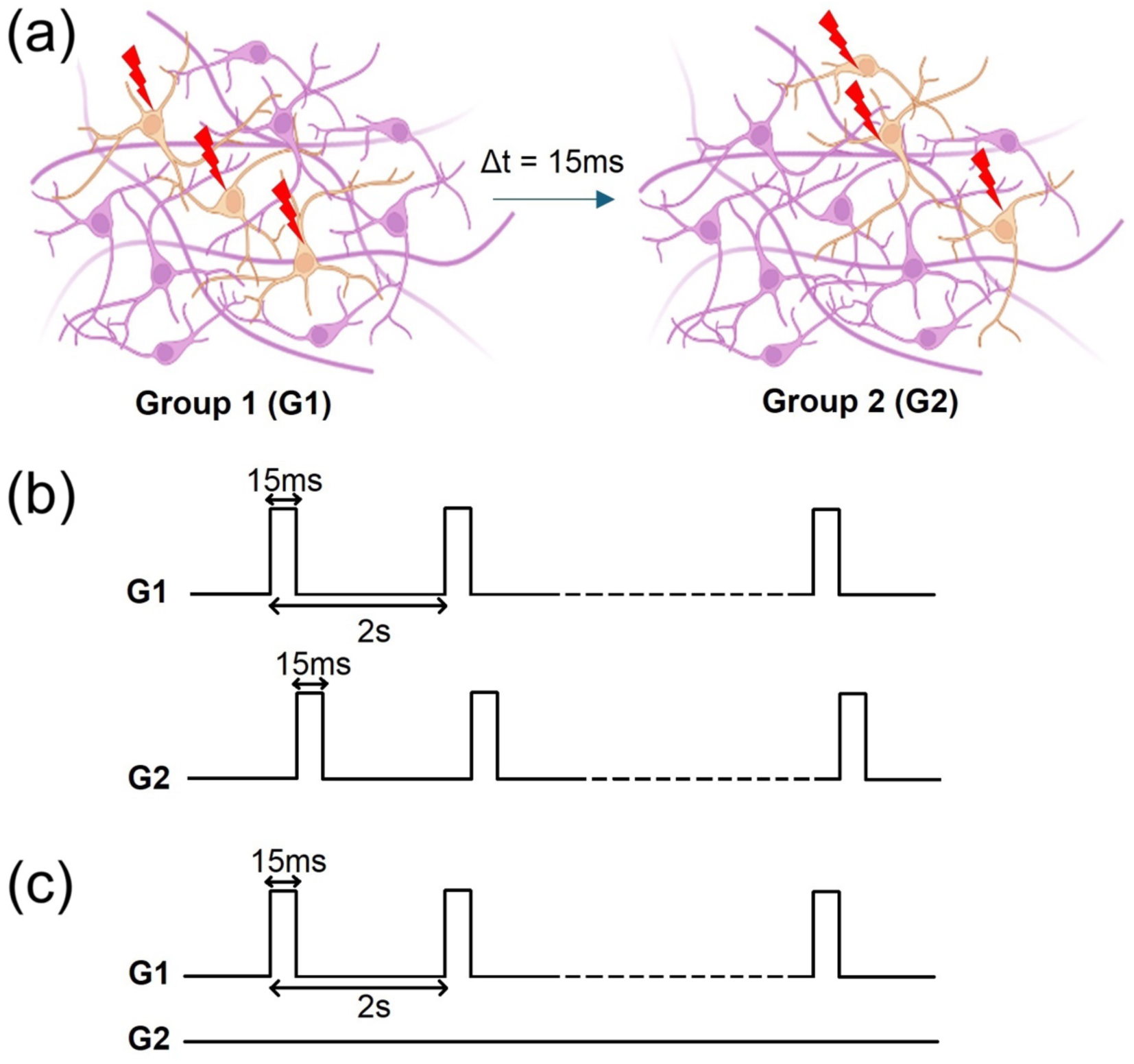
(a) Schematic of the training paradigm of two groups of neurons. The experiment is structured into three stages: training, testing 1, and testing 2. (b) **Training:** G1 neurons were first stimulated, followed by G2 neurons, using a pulse duration of 15 milliseconds. Stimulation occurred every 2 seconds for a total duration of 90 seconds. (c) **Testing:** Only G1 neurons were stimulated (20 times) to determine if G2 neurons would fire in response, indicating that G2 neurons had learned to fire after G1 neurons due to the training phase. A second testing phase involved stimulation of only G2.

## III. RESULTS

### A. Enhancing the reliability of training through background stimulation (context)

Background stimulation was systematically optimized to enhance the reliability of neuronal firing. During group-specific stimulation experiments, 10% of the background pixels within the DMD were utilized for stimulation, while pixels corresponding to the non-stimulated groups were excluded. A range of background stimulation levels (0%, 3%, 5%, 10%, 15%) were evaluated, with 10% identified as the optimal condition to achieve robust neuronal response rates. Prior to the experiment, global stimulation was applied to identify neurons that reliably responded to stimulation, as indicated by a 100% response rate. From this subgroup, 10 neurons were selected and designated as directly stimulated neurons (G1 and G2) for subsequent experiments.

Fig. 2 depicts the neuronal responses, inferred from ΔF/F calcium activity, for directly targeted neurons and neighboring neurons that responded to background stimulation. These neighboring neurons were identified based on their high correlation with the directly targeted neurons during the 10% background stimulation period. An offset of 0.35 was applied to each ΔF/F trace to improve visualization. In this experiment, a total of 10 neurons were directly targeted. For each background stimulation level evaluated, 20 stimulation pulses were delivered over a total duration of 40 seconds. Stimulation was performed using the DMD, which was controlled in real-time through the NeuroART software.

**Fig. 2.**
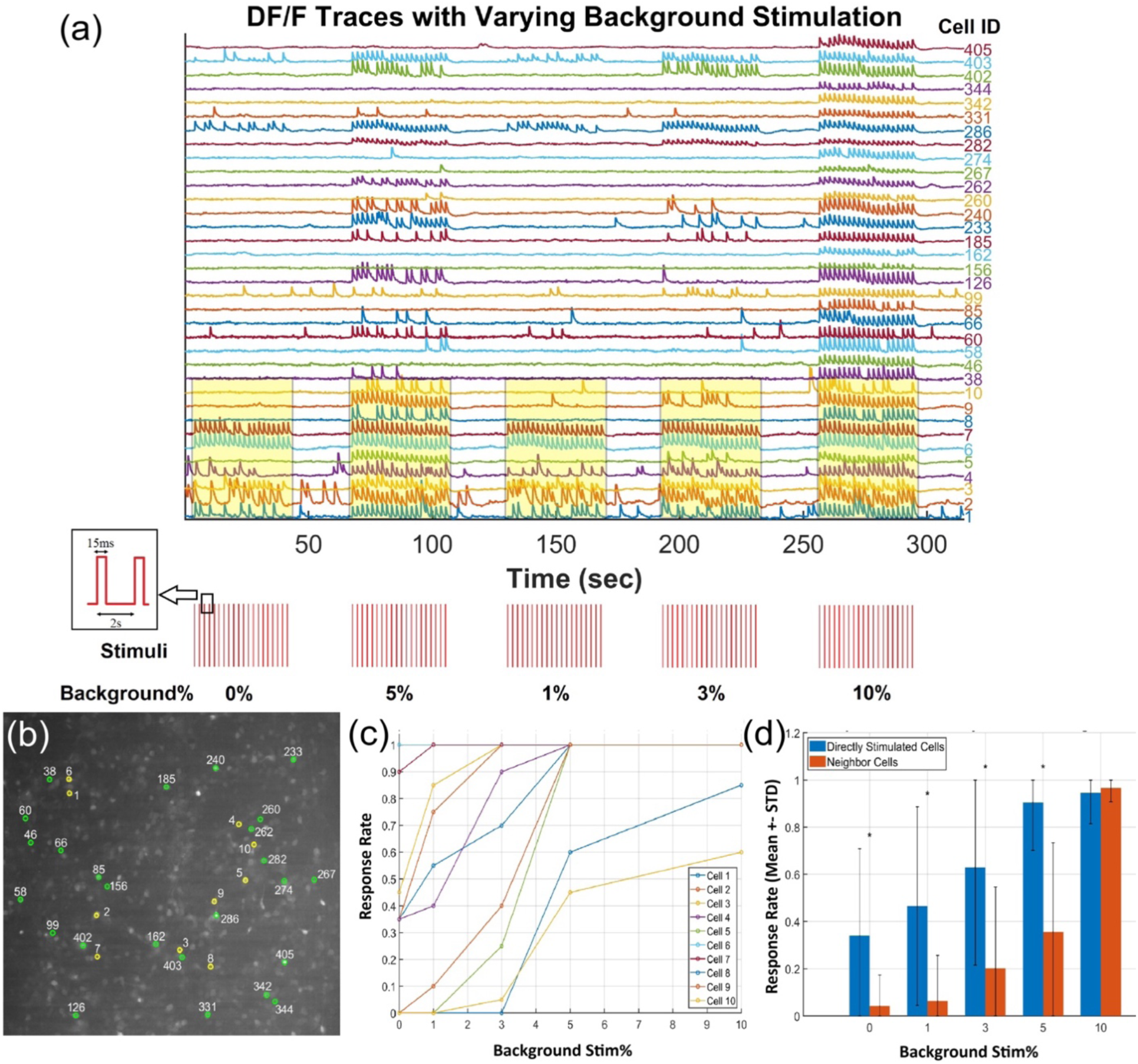
**(a) (Top)** ΔF/F traces of directly targeted neurons and neighboring cells responding to background stimulation. Directly targeted neurons are highlighted in yellow, and ΔF/F calcium activity traces are plotted with an offset of 0.35 to enhance visualization. The total number of cells in the field of view was 493. **(Bottom)** Timing of optogenetic stimulation (555 nm), delivered every 2 seconds as 15ms pulses using the DMD setup. Stimulation was provided for a total duration of 40 seconds (20 pulses) for each background stimulation level. The inset on the bottom left illustrates the pulse duration and the interval between consecutive pulses. **(b)** Spatial distribution of directly targeted neurons (yellow) and neighboring cells that reliably respond to 10% background stimulation (green). **(c)** Response rates of directly targeted neurons across different levels of background stimulation. The response rates of all targeted neurons show a monotonic increase with rising background stimulation levels. **(d)** Comparison of response rates between directly targeted neurons and neighboring cells, focusing only on those that reliably respond to 10% background stimulation.

The mean image of the imaging field of view indicates the spatial distribution of directly targeted neurons and reliably responding neighboring neurons in yellow and green, respectively (Fig. 2(b)). It was observed that for each directly targeted neuron, the response rate increased monotonically as the background stimulation level was raised from 0% to 10%. At 10% background stimulation, 8 out of the 10 directly targeted neurons achieved a 100% response rate (Fig. 2(c)). Additionally, 25 out of the 493 neurons within the field of view reliably responded to 10% background stimulation. These neighboring neurons may contribute contextual input that facilitates the reliable responses of the directly targeted neurons, as evidenced by the near-complete response rates observed at 10% background stimulation. Furthermore, at background stimulation levels below 10%, the response rates of directly targeted neurons were significantly higher than those of neighboring neurons that reliably responded to 10% background stimulation (Fig. 2(d)).

### B. Impact of pulse duration on the training paradigm and reliability of photostimulation

An additional factor considered was the pulse duration applied during the photostimulation of each neuronal group. Three experiments were conducted to evaluate the effect of stimulation pulse duration (10ms, 15ms, and 20ms) on group-to-group learning process in primary rat embryonic cortical neurons. All experiments (experiments 1–3) were performed using DIV10 neurons transduced as described in section II.A. For this analysis, only the directly targeted neurons in the training phase (i.e., neurons in G1 and G2) were considered.

As shown in Fig. 3, increasing the pulse duration from 10ms to either 15ms or 20ms resulted in a statistically significant increase in the response rate of the targeted neurons. Although increasing the pulse duration from 15ms to 20ms produced a slight increase in the overall response rate, this change was not statistically significant. To ensure that neurons learn according to Hebbian learning principles and spike timing-dependent plasticity (STDP) rules, it is critical to maintain the time delay between the activation of G1 and G2 below 20ms, as delays beyond this threshold may impede long-term plasticity. Therefore, a pulse duration of 15ms was identified as a reasonably suitable condition for training and testing group-to-group Hebbian learning, accounting for the variability in neuronal responsiveness to optogenetic stimulation.

**Fig. 3.**
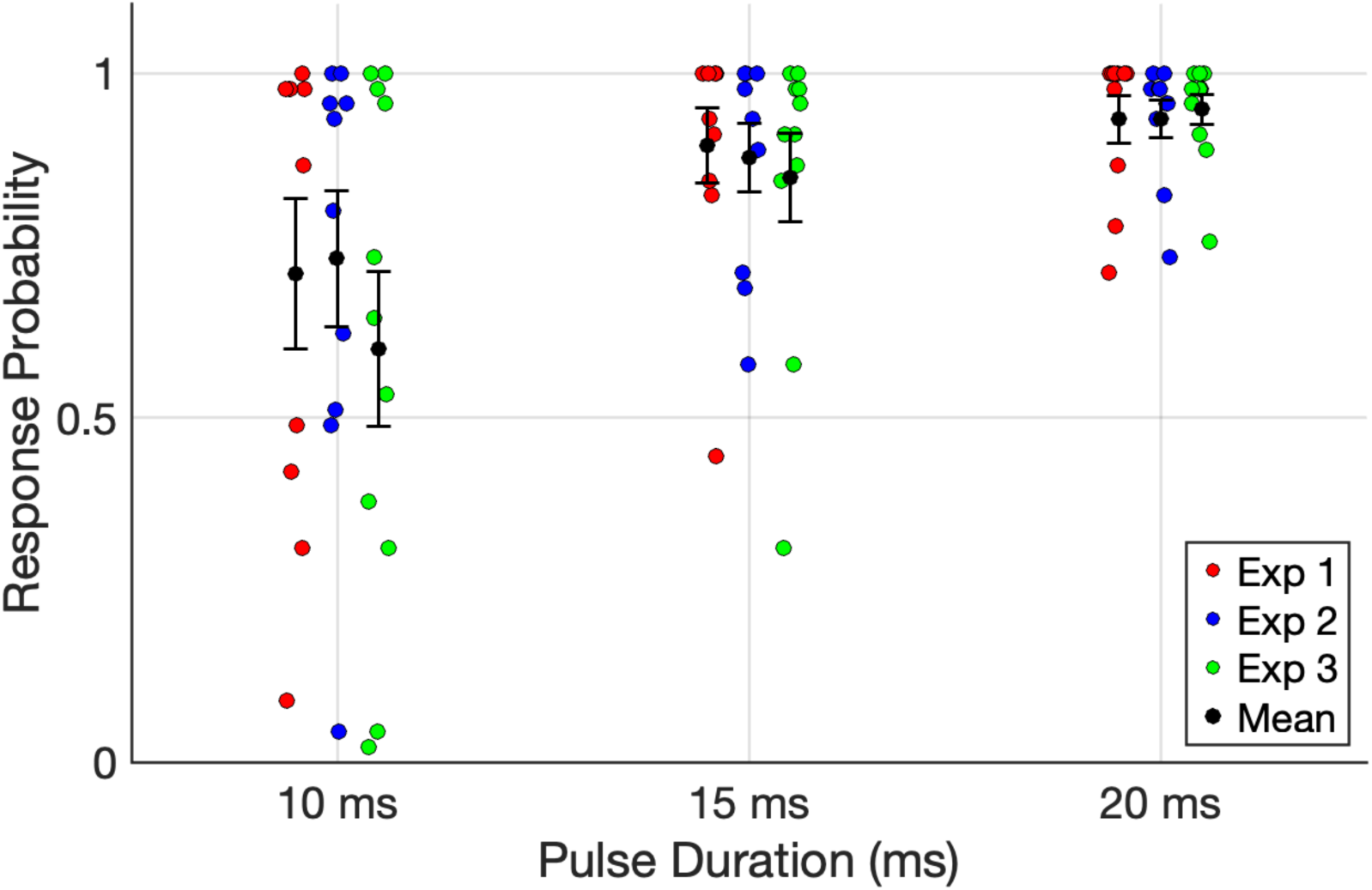
The impact of pulse duration on neuronal response probability. Three pulse durations (10ms, 15ms, and 20ms) were evaluated in each experiment. Red, blue, and green dots represent the response probabilities for the three experiments, while black dots indicate the mean response probability for each condition. Error bars represent the standard error of the mean (SEM). A slight horizontal offset and jitter were added to the scatter plots to improve clarity and avoid overlap of individual data points. The 10ms and 15ms response probabilities were significantly different (Wilcoxon Signed-Rank Test, included in the Appendix).

Considering the reliability of neuronal responses and the relevant timescales for Hebbian learning, as discussed in sections III.A and III.B, subsequent experiments investigating group-to-group Hebbian learning of neuronal groups were conducted using 15ms photostimulation pulses combined with 10% background stimulation.

### C. Training and testing of group-to-group Hebbian learning

As described in section II.B, 10 neurons were selected for the group-to-group Hebbian learning experiments. In these experiments, two groups of neurons, G1 and G2, each comprising 5 neurons, were trained by sequential stimulation. G1 was stimulated first with a 15ms pulse, followed by G2 with another 15ms pulse. A total of 45 pulses were delivered to each group over a training period of 90 seconds. To provide contextual input, a 10% background stimulation was applied in conjunction with the targeted stimulation. However, during the stimulation of one group, all pixels corresponding to the other group were excluded from the background stimulation to prevent unintended activation. The DMD, controlled in real-time using the NeuroART software, was employed for precise stimulation.

After the training phase, two rounds of testing were conducted to assess group learning and investigate the unidirectional nature of group-to-group Hebbian learning. During testing phase 1, only the neurons in G1 were stimulated. Notably, some neurons in G2 responded, indicating that learning had occurred after just 45 stimuli (within less than 2 minutes), benefiting from the collective reinforcement of shared synaptic activity, unlike individual pairwise training. In testing phase 2, when only G2 neurons were stimulated, there was no corresponding response from G1 neurons, confirming the unidirectional nature of the learned association. Fig. 4 illustrates the neuronal responses, inferred from ΔF/F calcium activity, for both directly targeted neurons and neighboring neurons that responded to the background stimulation. Neighboring neurons were identified based on their high correlation with the directly targeted neurons during the training phase with 10% background stimulation. For visualization clarity, an offset of 0.35 was applied to each ΔF/F trace. G1 neurons are highlighted in yellow, and G2 neurons are highlighted in light purple.

**Fig. 4.**
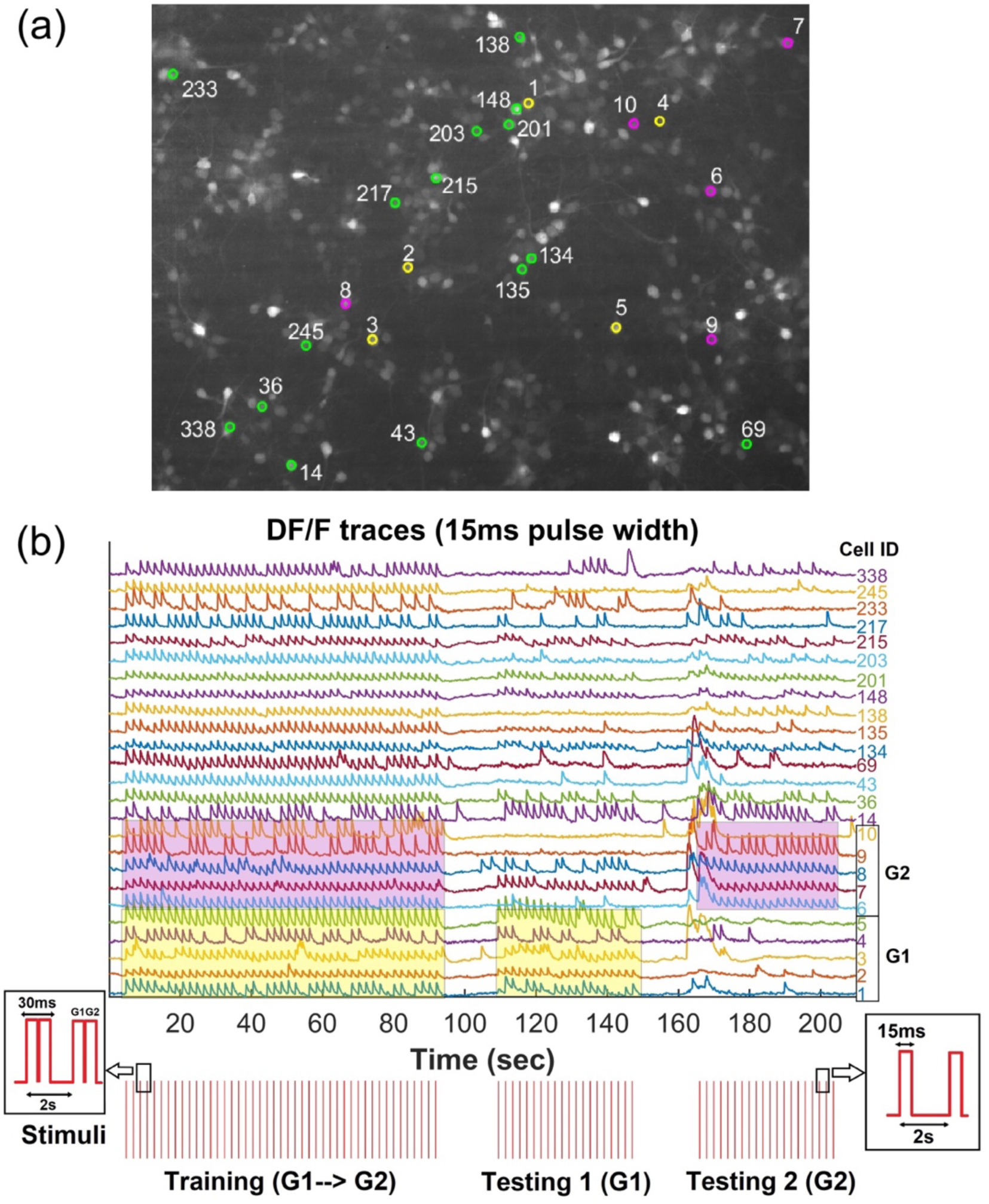
**(a)** Spatial distribution of directly targeted neurons in G1 (yellow), G2 (magenta) and neighboring cells that reliably responded to 10% background stimulation (green). **(b) (Top)** ΔF/F traces of directly targeted neurons and neighboring cells responding to background stimulation. Traces of directly targeted neurons are highlighted in yellow (G1) and light blue (G2). ΔF/F calcium activity traces are displayed with an offset of 0.35 for improved visualization. The total number of cells in the field of view was 361. **(Bottom)** Timing of optogenetic stimulation (555 nm), delivered every 2 seconds as 15ms pulses using the DMD setup. Stimulation was applied for a total duration of 90 seconds (45 pulses) during the training phase and 40 seconds (20 pulses) during each testing phase. The inset on the bottom left shows the pulse duration and intervals between consecutive pulses during training, while the inset on the bottom right shows the corresponding details for testing.

As illustrated in Fig. 5(a), almost all cells included in this analysis responded reliably during the training phase, where G1 and G2 neurons were directly photostimulated with a 15ms time delay, a timescale relevant for spike-timing-dependent plasticity (STDP)-based Hebbian learning. During the Testing 1 phase, only G1 neurons were directly stimulated, with the regions corresponding to G2 neurons excluded from background stimulation. Under these conditions, 2 out of 5 G2 neurons exhibited strong responses (response rates above 50%) by spiking in response to G1 activation. In contrast, no such responses were observed during the Testing 2 phase, where only G2 neurons were directly stimulated, confirming that the learning occurred unidirectionally from G1 to G2, with no reverse learning. Additionally, reliably responding neighboring neurons during the training phase demonstrated altered behavior during the testing phases, showing reduced response rates to background photostimulation, with some neurons responding to only a minimal number of stimuli. Notably, certain neighboring neurons (IDs: 135, 138, 203, 233, and 338) responded specifically during either the Testing 1 or Testing 2 phases, further supporting the cooperative role of neighboring neurons in providing contextual input to the learning process. Neighboring cells were further clustered based on their response rates during the Testing 1 phase to highlight patterns of contextual learning and variability in responses. The heatmap emphasizes the differences in neuronal responses across conditions, with higher response rates shown in red and lower response rates in blue.

**Fig. 5.**
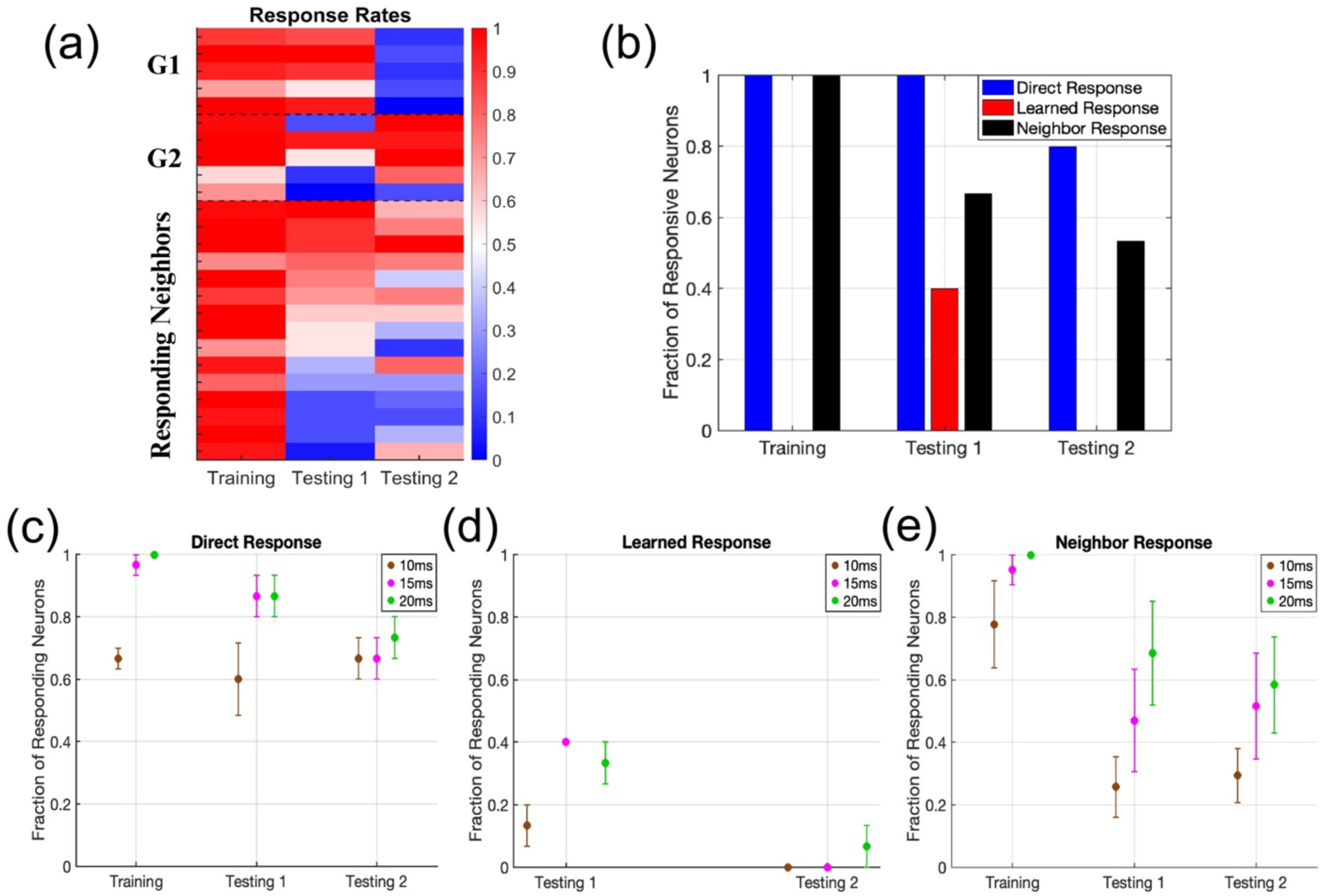
**(a)** Response probabilities for all analyzed cells during the phases, training, testing 1, and testing 2. Response probability ranges from 0 to 1, where 0 indicates no response and 1 indicates a 100% response to each photostimulation pulse. Cells are arranged by their group assignments: Group 1 (G1), Group 2 (G2), and neighboring cells. Black dotted lines separate G1, G2, and neighboring cells for clarity. **(b)** Bar graph illustrating the fraction of responsive neurons across experimental stages (Training, Testing 1, Testing 2) categorized by neuronal groups. Neurons are grouped as directly stimulated (blue), non-stimulated (red), and responding neighboring neurons (yellow). A neuron is classified as responsive if it generates a spike corresponding to each stimulus at least 50% of the time. Training includes only directly stimulated and responding neighboring neurons, while Testing 1 and Testing 2 include all three groups. **(c)** The mean fraction of responsive neurons among those directly stimulated during the training, testing 1, and testing 2 phases. **(d)** The mean fraction of responsive neurons among non-stimulated neurons that were trained during the training phase to respond following the activation of directly stimulated neurons (i.e., learned response). **(e)** The mean fraction of responsive neighboring neurons that consistently responded to a 20ms stimulation pulse duration during the training phase. Error bars represent the standard error of the mean (SEM) for pulse durations of 10ms, 15ms, and 20ms across each experimental stage.

Fig. 5(b) shows the fractions of responsive neurons across different categories: directly stimulated, non-stimulated, and responding neighbors. Neurons were classified as responsive if they responded to at least 50% of the stimuli. Hebbian learning was evident in 2 out of the 5 neurons in G2, which exhibited responses exceeding the 50% threshold. To further validate group-to-group Hebbian learning, the experiment was repeated on three different cell cultures at DIV10, utilizing three distinct stimulation pulse durations to investigate response variations. Fig. 5(c), (d), and (e) highlight the differences in the fractions of responsive and the impact of pulse duration on overall learning efficiency. Individual datapoints are provided in the appendix (a slight horizontal offset and jitter were applied to the scatter plots to enhance clarity and prevent overlap of individual data points). Notably, G2 neurons demonstrated learning, as evidenced by their consistent responses (greater than 50%) during the Testing 1 stage. Additionally, the higher learning rates observed with 15ms and 20ms pulse durations, compared to 10ms, underscore the importance of using a sufficiently long pulse duration to facilitate effective training.

To further confirm the involvement of spike timing-dependent plasticity (STDP) rules in group-to-group Hebbian learning, the time difference (ΔT) between spikes was estimated for each neuronal pair during the training phase. Since some neurons did not respond with 100% reliability to stimulation pulses, the estimation was performed only when corresponding spikes were observed in both neurons for a given stimulation pulse. The time delays for each matching spike pair were calculated, and the average time delay was considered as the representative (ΔT) for each neuronal pair. Spike timings and ΔT were determined as illustrated in Fig. 6(a). The spike onset was defined as the time point when the ΔF/F trace reached 10% of the maximum ΔF/F for the given spike. For a given neuronal pair, the spike onset times were calculated for both neurons, and the difference in onset times was used to determine the time difference (ΔT) for that pair. This process was repeated for all possible pairs among directly targeted neurons (G1 and G2) and neighboring neurons that reliably responded to 10% background stimulation.

**Fig. 6.**
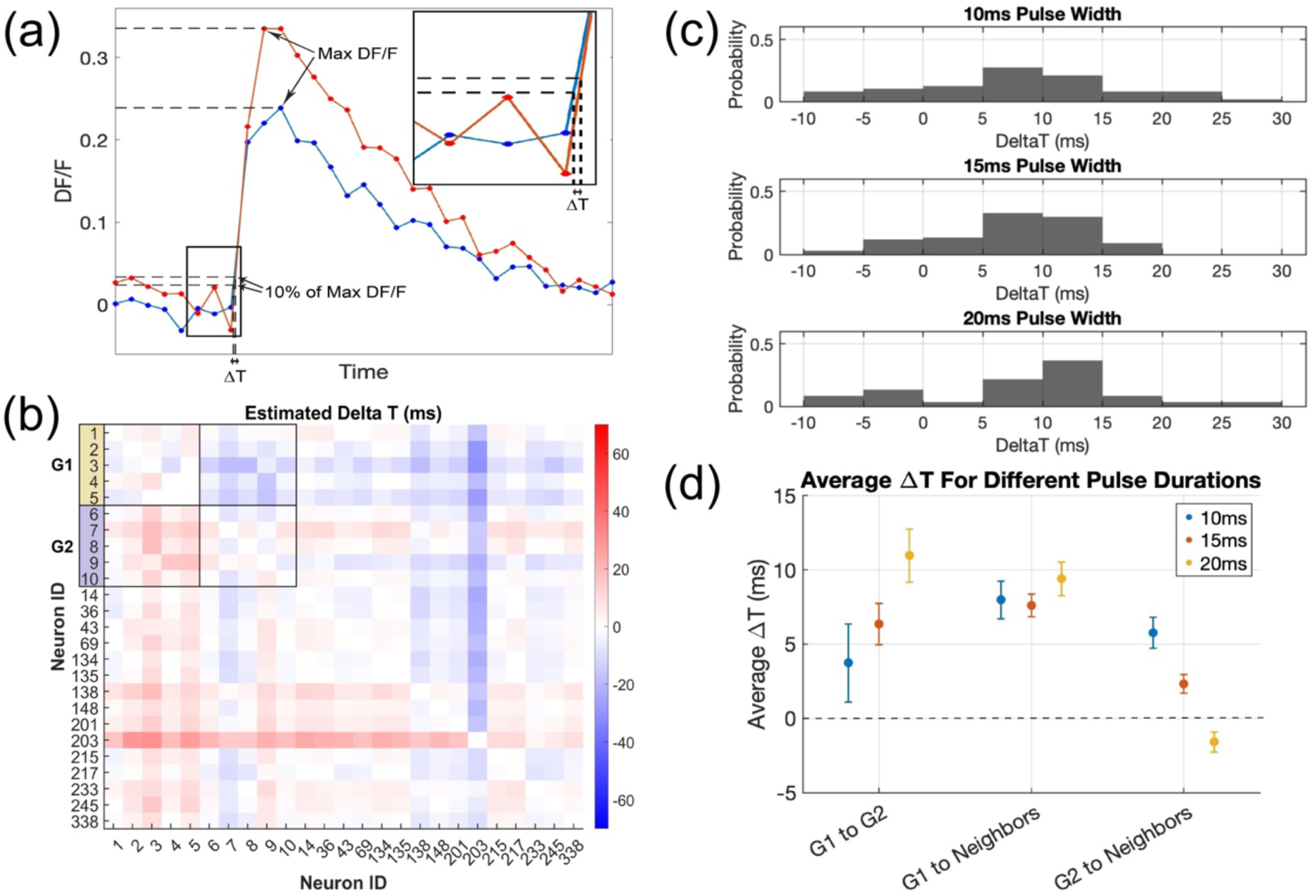
**(a)** A schematic of the spike timing and ΔT estimation based on the ΔF/F trace corresponding to each spike. The spike onset time for a given spike was identified as the time point at which the ΔF/F trace reached 10% of the maximum ΔF/F. ΔT is the difference between the spike onset times. The inset provides an enlarged view of the ΔT estimation. **(b)** Time delay (ΔT) matrix for all possible neuronal pairs, with red indicating positive ΔT (time advance) and blue indicating negative ΔT (time delay), read in row-to-column order. This analysis included directly stimulated neurons in G1 and G2, as well as neighboring neurons that reliably responded to 10% background stimulation. The top left corner (outlined using black squares) of the ΔT matrix corresponds to the directly targeted neurons (G1 and G2) **(c)** Histograms of ΔT between G1 and responding neighboring neurons, highlighting that most neighboring neurons are activated after G1 neurons during the training phase. **(d)** Mean ΔT across different groups, analyzed separately for each pulse duration. Error bars indicate the standard error of the mean (SEM).

Fig. 6(b) represents the ΔT matrix for all possible pairs, with red indicating positive ΔT (time advance) and blue indicating negative ΔT (time delay), read in row-to-column order. Notably, G2 neurons (cell IDs: 6 to 10) respond approximately 15ms after G1 neurons (cell IDs: 1 to 5). Additionally, neighboring neurons providing contextual input generally fired after G1 neurons, though not necessarily after G2 neurons. Overall, G2 neurons are activated after G1 neurons and the slight variability in time delays between G1 and G2 neurons could be attributed to the heterogeneity in calcium channel sensitivity to photostimulation light.

Fig. 6(c) presents histograms of ΔT values between G1 neurons and responding neighboring neurons for different stimulation pulse durations (10ms, 15ms, 20ms). The data emphasizes that most neighboring neurons are activated after G1 neurons during training, with ΔT values predominantly greater than zero. Fig. 6(d) illustrates the mean ΔT across different neuronal groups (G1 to G2, G1 to neighbors, and G2 to neighbors) for each pulse duration. Error bars represent the standard error of the mean (SEM). The results highlight a consistent trend of delayed activation in neighboring neurons, with longer pulse durations (15ms and 20ms) exhibiting more pronounced differences compared to shorter pulse durations (10ms). Notably, the mean ΔT between G1 and responding neighbors remains consistent and shows minimal variation with changes in pulse duration, suggesting a robust temporal relationship between these groups regardless of the stimulation parameters. At a pulse duration of 20ms, the mean ΔT between G2 neurons and their responding neighbors becomes negative, indicating that neighboring cells are activated before G2 neurons. This shift in activation timing may explain the bidirectional learning observed in some 20ms pulse duration experiments, as the earlier activation of neighboring cells could subsequently influence and drive the activity of G2 neurons after the training period.

## IV. DISCUSSION

Plasticity of neural networks plays a vital role in the development and integrative functions of the nervous system [2, 13-15]. The concept of STDP and its implications, such as persistent modifications of synaptic efficacy, has yielded new insights into the functional plasticity and dynamics of neural circuits. While pair-based STDP studies have provided critical foundational insights, limitations arise when extending these findings to network-level plasticity. Investigations into neuronal groups reveal additional modulatory complexities and neural basis of advanced cognitive functions, such as learning and memory, that are not observable in pairwise experimental setups [16]. Specifically, it would be beneficial for more studies to investigate cooperative effects of STDP within neuronal networks because most previous studies have not explored STDP beyond the scale of pairs of neurons.

This study provides insights into the mechanisms of group-to-group Hebbian learning, emphasizing the role of various factors such as pulse duration and background stimulation in effectively training neuronal networks. The results highlight key trade-offs and optimizations required for robust learning, along with the cooperative dynamics among neuronal subgroups and the contextual influence of neighboring neurons. One critical observation is the relative ease of training a group of neurons within a duration as short as 90 seconds compared to training individual neuronal pairs. Collective learning within groups appears to benefit from the collective reinforcement of shared synaptic activity, where interactions among multiple neurons contribute to enhanced plasticity. In contrast, training individual pairs may lack the collective reinforcement that facilitates stronger and more reliable learning. This finding aligns with the idea that neural circuits operate as functional units, where group-level interactions drive emergent properties, such as learning and memory. The collective plasticity observed in this study provides a framework for understanding how neuronal ensembles adapt and encode information efficiently.

Another critical finding of this study is the trade-off between pulse duration and the effectiveness of photostimulation. The results demonstrate that time delays induced by the pulse duration of photostimulation play a dual role: they must be sufficiently long to ensure effective stimulation but short enough to align with the timescales required for spike-timing-dependent plasticity (STDP). Longer delays, although allowing for robust stimulation, fail to induce learning due to their incompatibility with the constraints of STDP-based learning. Conversely, shorter delays, such as 10ms, do not provide enough stimulation to reliably activate the target neurons due to the heterogeneous sensitivities of neurons to photostimulation light. The 15ms pulse duration used for photostimulation represents a balance between these competing requirements, effectively driving STDP-based Hebbian learning while ensuring robust stimulation. This finding underscores the importance of timing in neuronal plasticity, particularly in the context of group-to-group learning, where precise temporal coordination is essential for directional learning and memory encoding.

The importance of context in the form of background stimulation was another key finding of this research. By incorporating 10% background stimulation, the reliability of neuronal responses was significantly enhanced. This context likely simulates a more naturalistic environment, where neurons receive input from multiple sources, mimicking the complexity of real-world synaptic activity. The background stimulation provides a baseline level of activity, enabling cooperative effects among neurons and facilitating robust group-level plasticity. Interestingly, the influence of context was not limited to the training phase but also extended to the testing phases, where background stimulation was combined with direct photostimulation. During the Testing 1 phase, the exclusion of G2 neurons from background stimulation revealed a directional learning process, where G1 activation led to responses in a subset of G2 neurons. In contrast, no such responses were observed during the Testing 2 phase, where G2 neurons were directly stimulated without G1 activation. This unidirectional learning process confirms that Hebbian learning in this system follows STDP-based rules, with precise timing and input specificity being critical for directional synaptic strengthening.

With the optimized parameters of 15ms pulse duration and 10% background stimulation, this study achieved effective training of two neuronal groups within a short duration of 90 seconds. This efficiency is noteworthy, as it demonstrates that group-to-group Hebbian learning can occur rapidly under optimized conditions, aligning with timescales relevant for physiological plasticity. The unidirectional nature of the learning process, where responses were observed from G1 to G2 but not in the reverse direction, further confirms the fidelity of the STDP-based Hebbian learning rules. This directional specificity underscores the importance of precise temporal coordination in driving plasticity and suggests that group-level learning mechanisms are inherently directional.

The behavior of neighboring neurons during the different phases of the experiment further highlights their cooperative role in the learning process. During the training phase, these neurons responded reliably to background stimulation, providing a contextual framework for the group-to-group learning of G1 and G2. However, during the testing phases, their responses varied significantly, with lower response rates observed overall. Some neurons exhibited minimal responses to background stimulation, suggesting a reduced contextual influence during the testing phases. Interestingly, a subset of neighboring neurons responded specifically during either the Testing 1 or Testing 2 phases. This observation highlights the nuanced role of neighboring neurons in shaping the learning process, where their contextual input appears to support and reinforce the activity of directly stimulated groups. The selective responses of these neighboring neurons underscore the complex interplay between local network dynamics and global learning mechanisms.

The insights from this study on group-to-group Hebbian learning offer a framework for exploring the cooperative character of information flow and learning in living neural networks. In particular, here we demonstrate the importance of background context in enabling consistent responses to optogenetic stimulation patterns, and we show that this context also contributes to network-level plasticity. Our finding – that learning of groups can be rapidly achieved in-vitro using self-assembled neural networks – opens avenues to investigate memory formation and even decision-making processes in-vitro under carefully controlled conditions within well-defined contexts.

## ACKNOWLEDGMENTS

This work was supported by the Air Force Office of Scientific Research (AFOSR) in the United States under grant FA9550-22-1-0405, with instrumentation support provided through grant FA9550-23-1-0065.

## APPENDIX

**Fig. A1.**
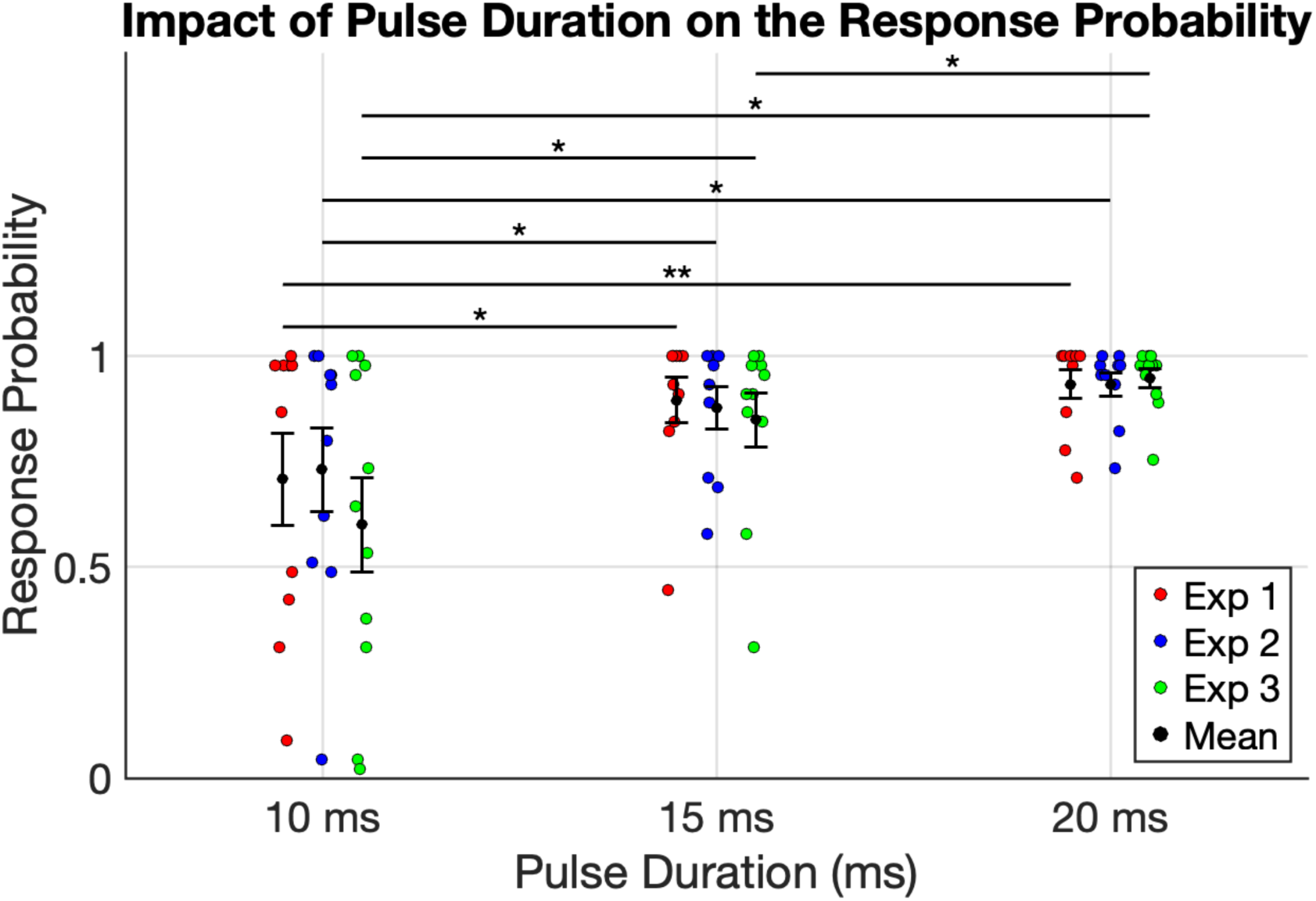
The effect of pulse duration on neuronal response probability. Three pulse durations (10ms, 15ms, and 20ms) were assessed in each experiment. Response probabilities from three experiments are represented by red, blue, and green dots, while black dots denote the mean response probability for each condition. Error bars indicate the standard error of the mean (SEM). A slight horizontal offset and jitter were applied to enhance clarity and minimize overlap of individual data points. Statistical analysis using the Wilcoxon Signed-Rank test revealed significant differences in response probabilities between the 10ms and 15ms pulse durations, as well as between the 10ms and 20ms pulse durations across all experiments.

**Fig. A2.**
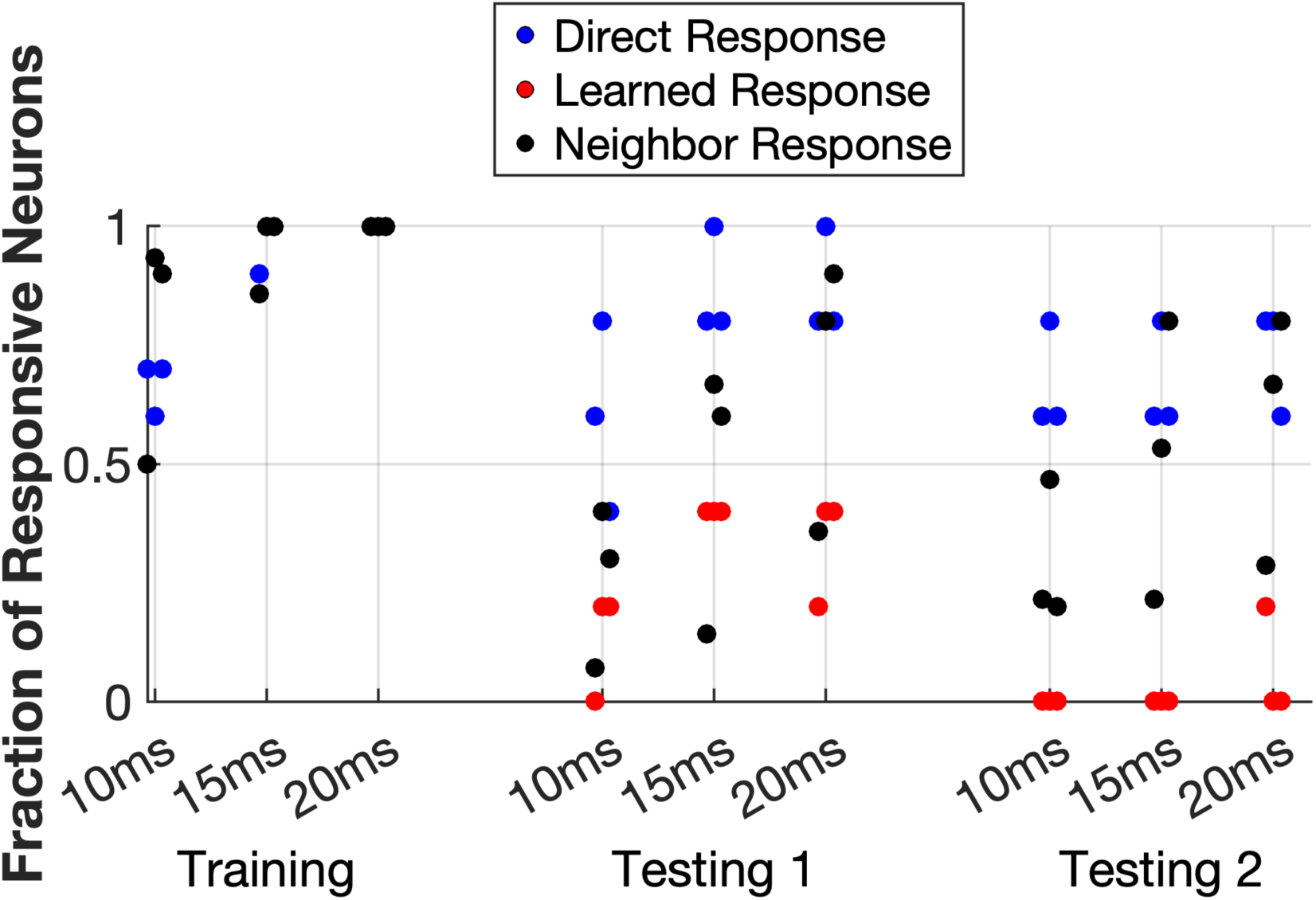
Fraction of responsive neurons (defined as those responding to at least 50% of stimuli) across different pulse durations (10ms, 15ms, and 20ms) during training and testing sessions in three biological replicates (imaged at DIV10). Three response types are depicted: direct responses (blue), learned responses (red), and neighbor responses (black). Learned responses originate from neurons that were not directly stimulated; G2 neurons during Testing 1 and G1 neurons during Testing 2. During training, all G1 and G2 neurons were targeted, and their responses were classified as direct. Each dot represents data from an individual experiment for a given condition. Learned responses are evident in Testing 1 but are less prominent in Testing 2, suggesting the unidirectional nature of learning. Neighboring neurons included in the analysis were those that exhibited consistent responses during training with a 20ms pulse duration. A slight horizontal jitter was applied to enhance visualization and to minimize data point overlap.

